# Layer 5 and 6b extratelencephalic neurons encode distinct sound features in auditory cortex

**DOI:** 10.64898/2026.03.19.712991

**Authors:** Madan Ghimire, Ross S. Williamson

**Affiliations:** Department of Otolaryngology, University of Pittsburgh, Pittsburgh, PA; Department of Neurobiology, University of Pittsburgh, Pittsburgh, PA; Department of Bioengineering, University of Pittsburgh, Pittsburgh, PA; Pittsburgh Hearing Research Center, University of Pittsburgh, Pittsburgh, PA; Center for the Neural Basis of Cognition, University of Pittsburgh, Pittsburgh, PA

## Abstract

Extratelencephalic (ET) neurons in layers (L)5 and 6b of the auditory cortex (ACtx) provide major corticofugal outputs to subcortical structures and contribute to auditory learning and experience-dependent plasticity. Although both ET subtypes innervate overlapping down-stream targets, they differ in morphology, intrinsic physiology, and molecular identity, suggesting distinct functional roles. However, their *in vivo* response properties remain incompletely characterized. Here, we used a projection-defined viral strategy to express GCaMP8s selectively in L5 and L6b ET neurons in mouse ACtx and recorded calcium activity during presentation of diverse acoustic stimuli. L5 ET neurons predominantly exhibited sound-evoked excitation, higher response sparseness, and greater trial-to-trial reliability than L6b ET neurons; these differences were most pronounced for pure tones and sinusoidally amplitude-modulated (sAM) noise. In contrast, L6b ET neurons frequently exhibited sound-evoked suppression, particularly for more complex stimuli (including sAM noise and spectrotemporal ripples). Unsupervised clustering identified separable temporal response motifs and tuning profiles. L5 ET neurons favored single-peaked frequency tuning and monotonic intensity tuning, whereas L6b ET neurons exhibited multi-peaked and complex frequency tuning, more non-monotonic intensity dependence, and stronger pairwise functional coupling (noise correlations). Together, these results support complementary corticofugal processing streams, with L5 ET neurons conveying more selective acoustic feature representations and L6b ET neurons conveying more integrative corticofugal signals.

## Introduction

Extratelencephalic (ET) neurons in the auditory cortex (ACtx) provide major descending corticofugal outputs to subcortical targets [1–3], projections that support auditory learning and experience-dependent plasticity [4–6]. ET neurons give rise to highly branched axons that broadcast auditory signals to multiple downstream structures, with the inferior colliculus (IC) of the midbrain serving as the most common target [7]. ET neurons also collateralize broadly within and beyond the telencephalon, innervating the medial geniculate body (MGB; auditory thalamus), striatum, amygdala, pons, and other nuclei [1, 2, 8–15].

ACtx ET neurons arise from distinct cortical layers, with the majority located in layer (L)5 and a smaller subset in L6b [11, 16–18]. Although both subtypes give rise to extensive collateral axons, their projection patterns and target preferences differ markedly. L5 ET neurons predominantly innervate midbrain structures (including the IC and superior colliculus (SC)), whereas L6b ET neurons preferentially target forebrain nuclei such as the MGB, striatum, and amygdala [10]. In addition to their distinct projection patterns, L5 and L6b ET neurons differ in morphology, intrinsic physiology (*in vitro*), and transcriptomic identity [12, 18–20]. L5 ET neurons typically exhibit a classic pyramidal morphology with a prominent, tufted apical dendrite extending to L1, positioning them to integrate thalamocortical, local intracortical, and long-range corticocortical inputs across cortical layers [18, 19]. In contrast, L6b ET neurons tend to have smaller, horizontally-oriented somata and thin dendrites that branch heavily near the soma and extend distally toward the pial surface [12]. These morphological differences likely contribute to distinct intrinsic electrophysiological properties, such as input resistance and firing pattern dynamics, as shown in *in vitro* recordings [12, 18]. At the transcriptomic level, many L5 ET neurons express Ctip2 and Fezf2 [20], whereas L6b ET neurons overlap substantially with FOXP2-expressing populations within L6 [10, 21, 22].

Given these laminar differences, one might expect L5 and L6b ET neurons to encode auditory information in functionally distinct ways. L5 ET neurons are often conceptualized as long-range “broadcast neurons” that exhibit broad, non-linear tuning to sound features, with limited involvement in local intracortical processing [3, 8, 23, 24]. By contrast, canonical descriptions of L6 corticofugal function derive largely from studies of layer 6a corticothalamic (CT) neurons. These neurons are typically characterized by sparse, narrowly tuned responses and a strong capacity to modulate local circuits via direct and cortico-thalamo-cortical feedback loops [8, 25, 26]. Deep L6 neurons, including L6b, receive prominent neuromodulatory inputs (e.g., cholinergic and dopaminergic) and express high levels of the corresponding receptor subtypes [22, 27–29], suggesting that state-dependent influences may differentially shape L6b ET activity relative to L5. Importantly, L6b ET neurons are morphologically and molecularly distinct from L6a CT neurons, yet their *in vivo* response properties remain largely uncharacterized.

These differences raise a central question: do L5 and L6b ET neurons convey largely overlapping auditory representations, or do they specialize in distinct aspects of sound processing? To address this, we performed two-photon calcium imaging in awake mice to monitor the activity of ACtx L5 and L6b ET neurons. To sample responses across a broad range of spectral and temporal modulations, we presented both simple pure tones and complex, modulated stimuli, including sinusoidally amplitude-modulated (sAM) noise and spectrotemporal ripples. We observed pronounced subtype-specific differences: the majority of sound-responsive L5 ET neurons were excited by sound, whereas L6b ET neurons frequently exhibited sound-evoked suppression. The subtypes also differed in frequency–intensity tuning selectivity and trial-to-trial response reliability, with differences most pronounced for pure tones. L5 and L6b ET neurons also differed in functional coupling, as measured by noise correlations. Collectively, these results suggest that L5 and L6b ET neurons participate in complementary corticofugal output streams that convey distinct aspects of auditory information.

## Results

To investigate functional differences between ACtx L5 and L6b ET neurons, we used a projection-defined viral strategy to selectively express GCaMP8s in corticocollicular neurons. Specifically, we injected a retrogradely transported virus encoding Cre recombinase (CAV2-Cre) into the right IC, followed by injection of a Cre-dependent AAV encoding GCaMP8s into ipsilateral ACtx, yielding expression in corticocollicular ET neurons spanning both L5 and L6b (**Figure 1A-D**). We localized primary ACtx using its tonotopic gradient revealed by widefield calcium imaging (**Figure 1E**), and assigned neurons as L5 or L6b based on somatic depth (L5: ∼ 400 − 550*µ*m; L6b: ∼ 700–830*µ*m below the pial surface) in separate two-photon imaging sessions targeting each layer. We then recorded neural activity in awake, head-fixed mice during passive listening to 50 ms pure tones (L5: *N* = 8 mice, *n* = 1280 neurons; L6b: *N* = 7, *n* = 689), 500 ms pure tones (L5: *N* = 3, *n* = 449; L6b: *N* = 3, *n* = 234), 500 ms sAM noise (L5: *N* = 9, *n* = 1567; L6b: *N* = 5, *n* = 622), and 500 ms spectrotemporal ripples (L5: *N* = 6, *n* = 766; L6b: *N* = 5, *n* = 451) (**Figure 1F-H**).

**Figure 1:**
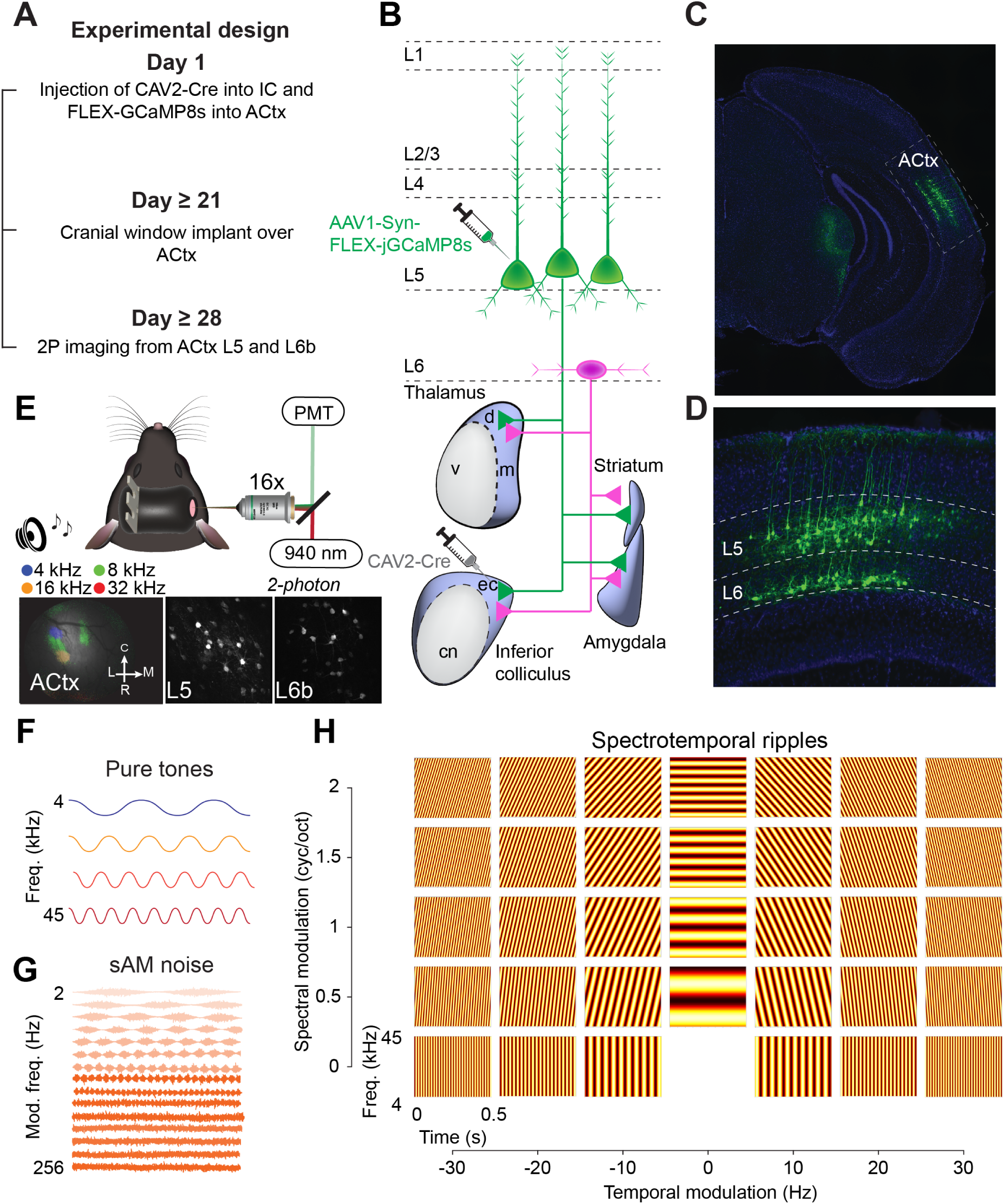
Experimental design and two-photon imaging of L5 and L6b ET neurons in ACtx. **A**: Experimental timeline showing virus injection, cranial window implantation, and two-photon imaging sessions. **B**: Schematic of the projection-defined viral strategy. A retrogradely transported virus encoding Cre recombinase (CAV2-Cre) was injected into the right inferior colliculus (IC), and a Cre-dependent AAV encoding GCaMP8s was injected into ipsilateral ACtx, yielding selective expression in corticocollicular ET neurons spanning L5 and L6b. **C**: Epifluorescence image showing GCaMP8s expression in ACtx. **D**: Magnified coronal section of ACtx showing GCaMP8s-expressing ET neurons in L5 and L6b. **E**: Top, schematic of the two-photon imaging configuration. Bottom left, example widefield epifluorescence image used to identify primary ACtx based on its tonotopic gradient. Bottom middle and right, example two-photon fields of view from L5 and L6b ET neurons, respectively, acquired at 2× zoom with a 16× water-immersion objective. **F-H**: Schematic representations of the three acoustic stimulus classes: pure tones **(F)**, sinusoidally amplitude-modulated (sAM) noise **(G)**, and spectrotemporal ripples **(H)**.

### L5 and L6b ET neurons exhibit distinct excitation and suppression profiles

ET neurons exhibited diverse sound-evoked response profiles, including both increases (excitation) and decreases (suppression) in activity (**Figure 2A-D**). We first quantified, for each stimulus class, the proportion of L5 and L6b ET neurons that were significantly excited or suppressed within a response window aligned to stimulus presentation (see Methods). Across mice and stimulus classes, the per-animal proportion of sound-excited neurons was ∼ 2 − 3 fold higher in L5 than in L6b (**Figure 2E**, *left*, Wilcoxon rank-sum test, *p* = 2.1 × 10^−5^), whereas L6b ET neurons showed a ∼ 3 − 4 fold higher incidence of sound-evoked suppression relative to L5 (**Figure 2E**, *middle*, Wilcoxon rank-sum test, *p* = 0.003). We next asked how these proportions varied across stimuli within each subtype. In L5, the fraction of excited neurons did not differ significantly across stimuli (**Figure 2F**, one-way ANOVA, *p* = 0.073). In L6b, the fraction of excited neurons increased with stimulus complexity, from tones to temporally and spectrotemporally modulated sounds (**Figure 2F**, one-way ANOVA, *p* = 0.013), with the largest difference between 50 ms pure tones and ripples (Tukey’s post-hoc test, *p* = 0.009). A similar divergence was observed for suppressed responses. In L5, the fraction of suppressed neurons did not vary significantly across stimuli (**Figure 2G**, one-way ANOVA, *p* = 0.311), whereas in L6b, suppression increased with stimulus complexity (**Figure 2G**, one-way ANOVA, *p* = 0.001), with the largest difference between 50 ms tones and sAM noise (Tukey’s post-hoc test, *p* = 9.0 × 10^−4^) and smaller differences between 50 ms tones and ripples (*p* = 0.032), and 500 ms tones and sAM noise (*p* = 0.020).

**Figure 2:**
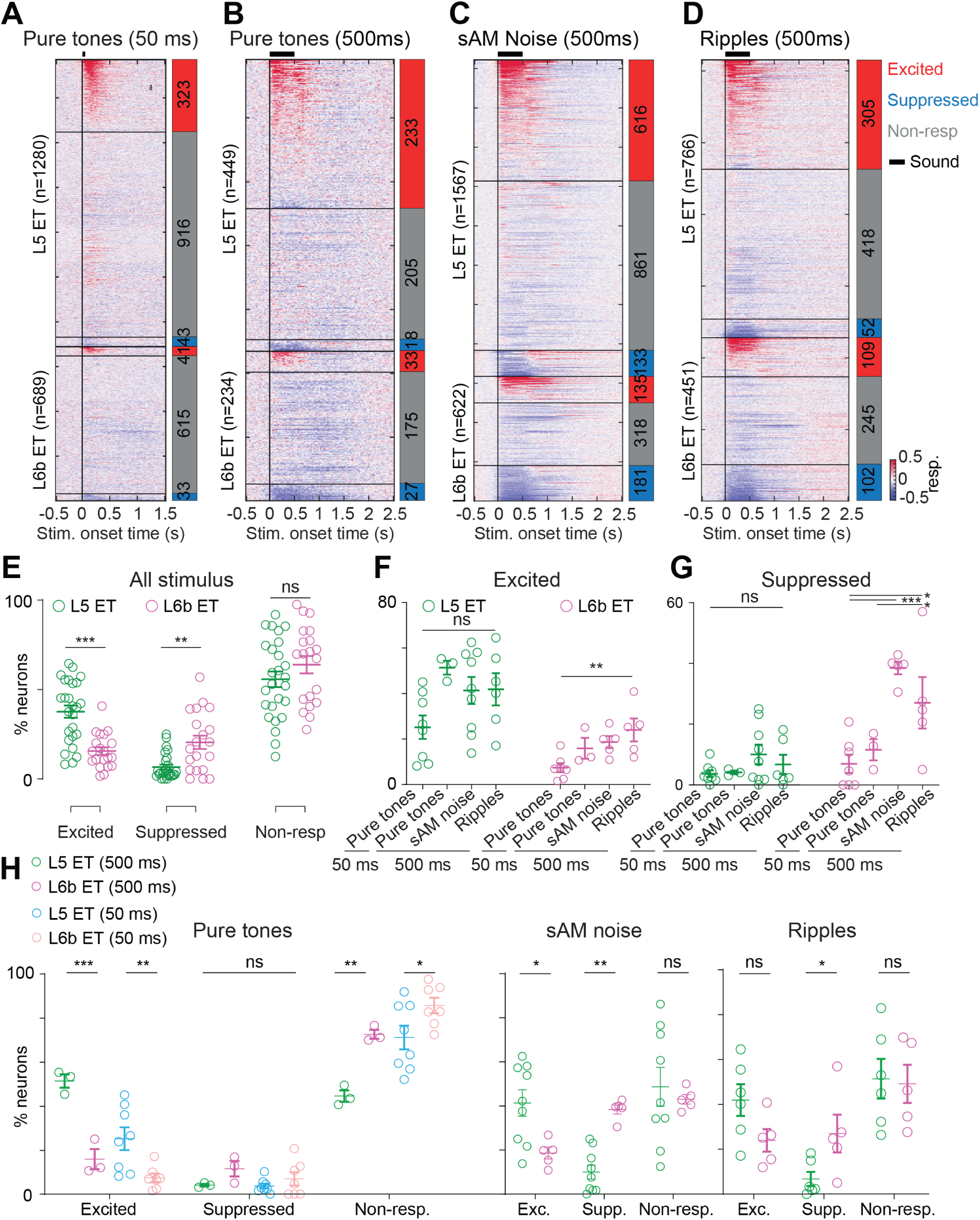
L5 and L6b ET neurons exhibit distinct sound-evoked response profiles. **A**: Mean peri-stimulus time histograms (PSTHs) for all L5 (top) and L6b (bottom) ET neurons in response to 50 ms pure tones, sorted by maximum response within the response window. Red bars indicate sound-excited neurons, blue bars indicate sound-suppressed neurons, and gray bars indicate non-responsive neurons. **B-D**: Same as A for 500 ms pure tones **(B)**, 500 ms sAM noise **(C)**, and 500 ms spectrotemporal ripples **(D)**. **E**: Per-animal proportion of sound-excited (left), sound-suppressed (middle), and non-responsive (right) neurons, pooled across all stimulus conditions. Each circle represents an individual animal. **F-G**: Per-animal proportion of sound-excited **(F)** and sound-suppressed **(G)** neurons for each stimulus condition, shown separately for L5 and L6b ET neurons. **H**: Proportion of L5 (green) and L6b (magenta) ET neurons in each response category (excited, suppressed, non-responsive) for 50 and 500 ms pure tones (left), 500 ms sAM noise (middle), and 500 ms spectrotemporal ripples (right). Asterisks denote statistically significant differences: Wilcoxon rank-sum test **(E)**, Tukey’s post-hoc test after one-way ANOVA **(F, G)**, and Šidák’s post-hoc test after two-way ANOVA **(H)**. * *p* ≤ 0.05, ** *p* ≤ 0.01, *** *p* ≤ 0.001, ns = non-significant.

Because L5 and L6b ET neurons showed distinct patterns of excitation and suppression across stimuli, we next asked whether, for each stimulus, the distribution of response types (excited, suppressed, non-responsive) differed between subtypes. Across all stimulus classes, there was a significant interaction between subtype and response type (**Figure 2H**, two-way ANOVA, subtype × response type interaction: 50 ms pure tones: *p* = 9.2 × 10^−10^, 500 ms pure tones: *p* = 6.2 × 10^−6^, sAM noise: *p* = 9.0 × 10^−4^, ripples: *p* = 0.040). Post hoc comparisons showed that L5 had a higher proportion of excited neurons in response to pure tones and sAM noise (**Figure 2H**, two-way ANOVA with Šidák’s post-hoc test, 50 ms pure tones: *p* = 0.003, 500 ms pure tones: *p* = 1.6 × 10^−4^, sAM noise: *p* = 0.044), whereas L6b had a higher proportion of suppressed neurons in response to sAM noise and ripples (**Figure 2H**, two-way ANOVA with Šidák’s post-hoc test, sAM noise: *p* = 0.009, ripples: *p* = 0.017). The proportion of non-responsive neurons was also higher in L6b during pure tone presentation (**Figure 2H**, two-way ANOVA with Šidák’s posthoc test, 50 ms pure tones: *p* = 0.022, 500 ms pure tones: *p* = 0.003). Together, these results indicate that excitation predominates among sound-responsive L5 ET neurons, whereas a larger fraction of L6b ET neurons show sound-evoked suppression. These patterns are consistent with L5 ET neurons receiving stronger net excitatory drive and L6b ET neurons being subject to more prominent inhibitory influences, in line with prior observations in deep-layer corticofugal circuits [25].

### L5 and L6b ET neurons exhibit diverse temporal response motifs across stimulus classes

Motivated by ET subtype differences in response profile (excitation versus suppression), we next asked whether their temporal response patterns differed. We observed a variety of temporally complex response profiles in both L5 and L6b ET neurons. To characterize these responses beyond simple excitation or suppression, we applied an unsupervised hierarchical clustering analysis to peri-stimulus time histograms (PSTHs) (**Figure 3A**; see Methods). For each stimulus class, we normalized PSTHs and clustered them separately. We then visualized the results by sorting neurons by cluster membership and plotting the ordered PSTHs (**Figure 3B-E** *left*), together with the corresponding cluster-mean profiles (**Figure 3B-E** *middle*). Finally, we quantified the proportions of L5 and L6b ET neurons in each cluster (**Figure 3B-E** *right*). This analysis revealed subpopulations with distinct temporal response motifs across stimulus classes.

**Figure 3:**
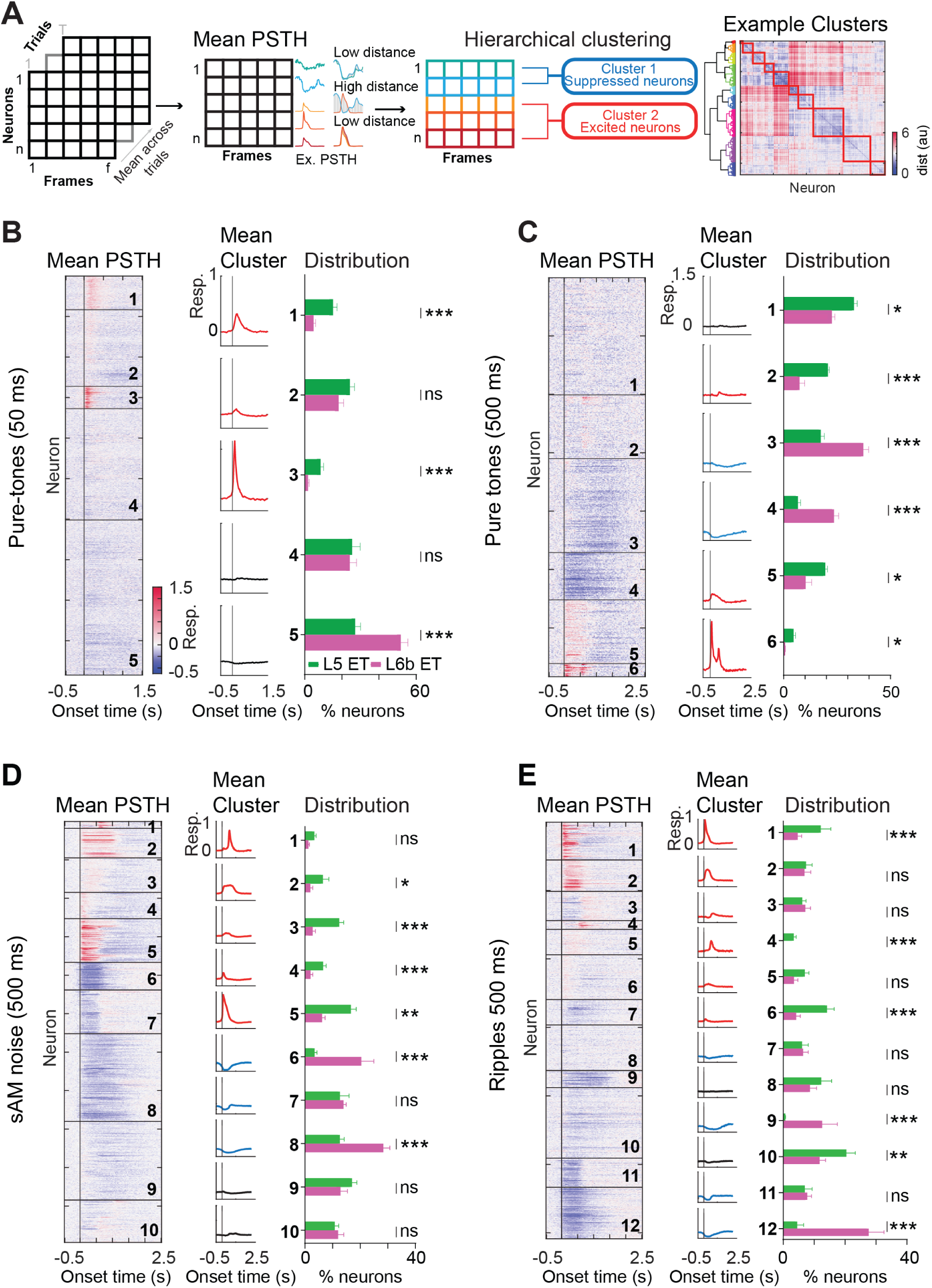
Stimulus-specific temporal response patterns differentiate L5 from L6b ET neurons. **A**: Schematic of the hierarchical clustering approach. Mean PSTHs were rescaled to the range [0, 1] and pairwise Euclidean distances were computed between all neurons. Right, example dendrogram with color-coded clusters and the corresponding distance matrix sorted by cluster membership. **B**: PSTHs for all neurons in response to 50-ms pure tones, organized by cluster identity (left), mean cluster response profiles (middle), and proportion of L5 and L6b ET neurons in each cluster (right). **C-E**: Same as B for 500 ms pure tones **(C)**, 500 ms sAM noise **(D)**, and 500 ms spectrotemporal ripples **(E)**. Asterisks denote statistically significant differences using chi-squared test with Bonferroni correction. * *p* ≤ 0.05, ** *p* ≤ 0.01, *** *p* ≤ 0.001, ns = non-significant.

For short pure tones (50 ms), most sound-evoked increases in activity were brief and dominated by onset responses (**Figure 3B**), consistent with the limited stimulus duration. In contrast, longer pure tones (500 ms) commonly drove sustained increases throughout the stimulus and, in a subset of neurons, clear offset responses (**Figure 3C**). Sound-evoked decreases in activity were prominent for 500 ms tones but comparatively rare for 50 ms tones (**Figure 3B-C**). This difference may reflect the progressive recruitment of sustained inhibitory mechanisms during longer stimuli, as cortical interneuron subtypes that mediate prolonged suppression, such as somatostatin-expressing neurons, operate on slower timescales than those mediating rapid onset inhibition [25, 30, 31]. Complex stimuli elicited a broader repertoire of temporal response patterns, including combinations of onset, sustained, and offset activity, as well as prolonged suppression (**Figure 3D-E**). Unsupervised clustering revealed ∼10-12 distinct temporal motifs per stimulus class (**Figure 3D-E**).

Cluster composition differed strongly by ET subtype for every stimulus class (chi-squared test, **Figure 3B** *right*, 50 ms pure tones: *p* = 1.9×10^−18^, **Figure 3C** *right*, 500 ms pure tones: *p* = 7.01× 10^−18^, **Figure 3D** *right*, 500 ms sAM noise: *p* = 1.3 × 10^−54^ and **Figure 3E** *right*, 500 ms ripples: *p* = 1.5×10^−43^). Pairwise comparisons showed that excited motifs were predominantly composed of L5 ET neurons (chi-squared test with Bonferroni correction, 50 ms pure tones: **Figure 3B**, clusters 1-3, *p* = 7.1 × 10^−8^, 0.075, 7.8 × 10^−7^; 500 ms pure tones: **Figure 3C**, clusters 5-6, *p* = 0.039, 0.017; 500 ms sAM noise: **Figure 3D**, clusters 1-5, *p* = 0.970, 0.040, 5.9 × 10^−6^, 1.3 × 10^−6^, 3.8 × 10^−4^; 500 ms ripples: **Figure 3E**, clusters 1-6, *p* = 3.9 × 10^−4^, 0.999, 0.999, 9.0 × 10^−4^, 0.230, 5.4 × 10^−7^), whereas suppressive motifs were largely dominated by L6b ET neurons (chi-squared test with Bonferroni correction, 500 ms pure tones: **Figure 3C**, clusters 3-4, *p* = 8.7 × 10^−9^, 1.7 × 10^−7^; 500 ms sAM noise: **Figure 3D**, clusters 6-8, *p* = 1.0 × 10^−15^, 0.999, 1.0 × 10^−15^; 500 ms ripples: **Figure 3E**, clusters 9, 11, 12, *p* = 2.9 × 10^−14^, 0.026, 1.0 × 10^−15^). Together, these results indicate that sound-responsive L5 ET neurons span a broad, heterogeneous repertoire of temporal motifs, while L6b ET neurons are over-represented among prolonged suppression motifs, particularly for longer and more complex stimuli.

### L5 and L6b ET neurons exhibit distinct frequency and intensity tuning profiles

Having established that L5 and L6b ET neurons differ in their response profiles, we next examined their frequency response areas (FRAs) to determine whether the two subtypes also differ in tuning to specific acoustic features. For each neuron with a significant excitatory pure tone response, we computed its best frequency and best intensity as the stimulus that evoked the strongest mean response when averaged across all intensities or frequencies, respectively (**Figure 4A**). The resulting distributions revealed significant tuning differences between L5 and L6b ET neurons. L5 ET neurons tended to prefer higher-intensity and lower-frequency tones, whereas L6b ET neurons responded more strongly to lower-intensity sounds and exhibited a broader distribution of best frequencies (**Figure 4B**, chi-squared test, best intensity: *p* = 1.4 × 10^−7^; **Figure 4C**, best frequency: *p* = 0.001).

**Figure 4:**
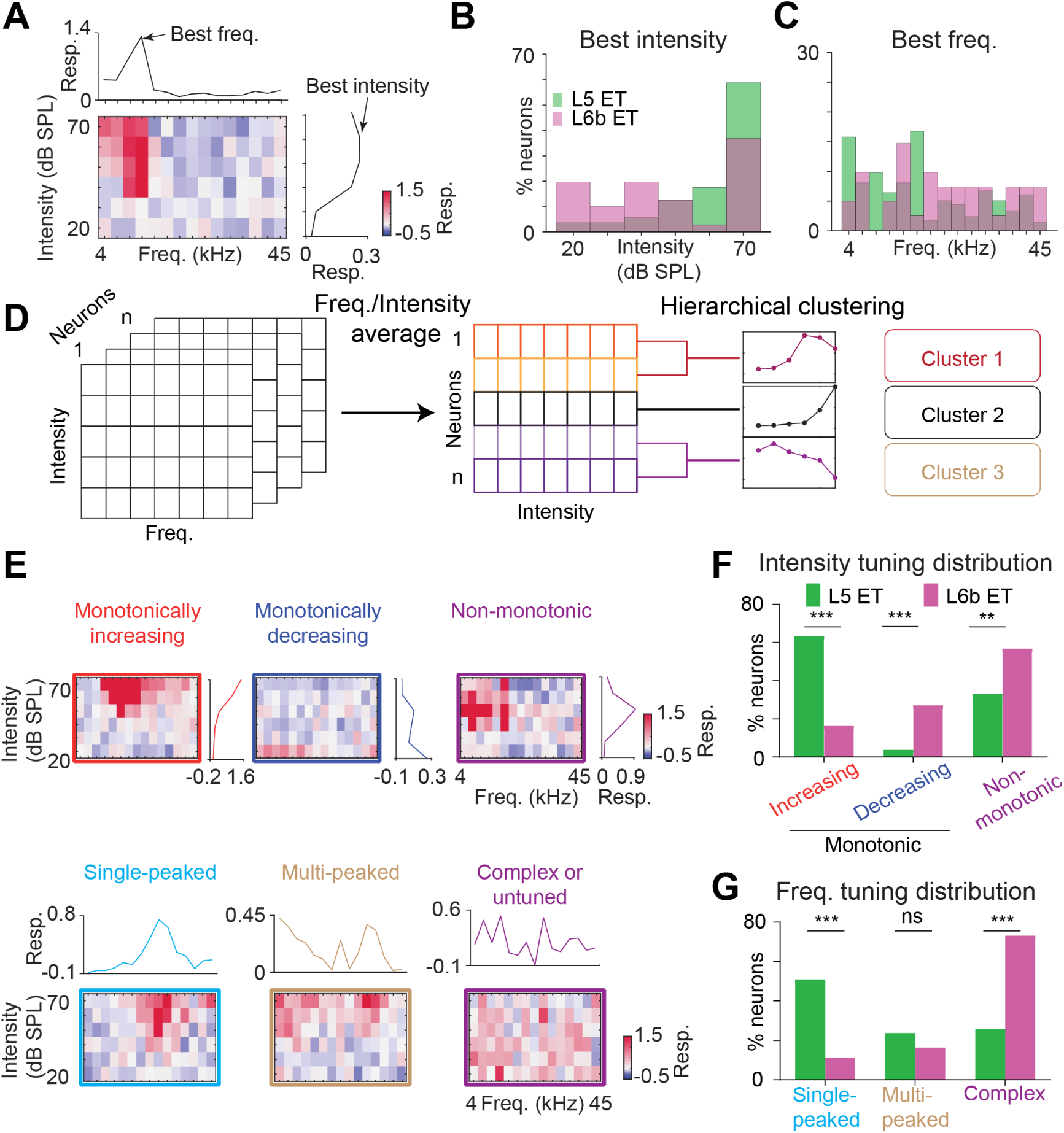
Pure tone tuning differs between L5 and L6b ET neurons. **A**: Example frequency response area (FRA) with frequency on the x-axis and intensity on the y-axis. Top, intensity-averaged response used to identify best frequency (peak). Right, frequency-averaged response used to identify best intensity (peak). **B-C**: Distribution of best intensity **(B)** and best frequency **(C)** for L5 and L6b ET neurons. **D**: Schematic of hierarchical clustering of pure tone frequency and intensity tuning profiles. Right, example dendrogram showing clustered tuning profiles, which were subsequently reclassified as monotonically increasing, monotonically decreasing, or non-monotonic (intensity) and single-peaked, multi-peaked, or complex (frequency). **E**: Top, example FRAs depicting monotonically increasing (left), monotonically decreasing (middle), and non-monotonic (right) intensity tuning profiles. Bottom, example FRAs depicting single-peaked (left), multi-peaked (middle), and complex (right) frequency tuning profiles. **F-G**: Distribution of intensity **(F)** and frequency **(G)** tuning categories for sound-excited L5 and L6b ET neurons. Asterisks denote statistically significant differences using chi-squared test with Bonferroni correction. * *p* ≤ 0.05, ** *p* ≤ 0.01, *** *p* ≤ 0.001, ns = non-significant.

We then performed unsupervised clustering of all frequency and intensity tuning profiles to identify distinct tuning motifs (**Figure 4D**). Intensity tuning curves were categorized as monotonically increasing, monotonically decreasing, or non-monotonic (**Figure 4E** *top*), and frequency tuning curves as single-peaked, multi-peaked, or complex (**Figure 4E** *bottom*). Most L5 ET neurons exhibited monotonic intensity tuning, reflecting higher pure tone thresholds, whereas L6b ET neurons more often showed non-monotonic profiles, with peak activity at intermediate intensities or decreasing activity at higher intensities (**Figure 4F**, chi-squared test with Bonferroni correction, monotonically increasing: *p* = 3.0 × 10^−11^, monotonically decreasing: *p* = 1.4 × 10^−5^ and non-monotonic: *p* = 0.002). L5 ET neurons were more likely to exhibit single-peaked tuning, whereas L6b ET neurons predominantly exhibited multi-peaked and complex tuning (**Figure 4G**, chi-squared test with Bonferroni correction, single-peaked: *p* = 2.9×10^−9^, multi-peaked: *p* = 0.574, complex: *p* = 6.9 × 10^−11^). This difference in tuning category distribution indicates that L6b ET neurons respond to a less restricted set of pure-tone combinations. Similar multi-peaked tuning has been reported in ACtx deep-layer subplate neurons [32], suggesting that such response patterns may reflect shared circuit or developmental origins of deep-layer neuronal populations. Together, these results support a model in which L5 ET neurons receive stronger or more selective thalamocortical input, yielding predominantly monotonic tuning to individual frequency channels [13]. In contrast, the prevalence of non-monotonic intensity tuning in L6b ET neurons is consistent with imbalanced excitatory and inhibitory synaptic inputs, where inhibition grows disproportionately with stimulus intensity, as has been demonstrated in non-monotonic neurons elsewhere in ACtx [33, 34]. The broader, multi-peaked frequency tuning of L6b ET neurons may additionally reflect convergent intracortical input [25, 35].

### L5 and L6b ET neurons exhibit similar tuning to sAM noise

To probe sensitivity to temporal envelope structure beyond simple tonal features, we presented sAM noise with varying modulation frequency and depth to assess amplitude-modulation processing, which supports speech envelope perception [36–38]. Previous work has shown that cortical neurons encode modulation features, such as modulation frequency and depth [39, 40]. Accordingly, we calculated the best modulation depth and best modulation frequency, defined as the stimulus that elicited the strongest responses averaged across modulation frequency or depth, respectively (**Figure 5A**). We found that L5 ET and L6b ET neurons were tuned to modulation frequencies spanning 2-256 Hz (**Figure 5B-C**). In contrast to pure tone responses, the distributions of best modulation frequency were largely similar between L5 ET and L6b ET neurons, but the distributions of best modulation depth differed (**Figure 5B-C**, chi-squared test, best modulation depth: *p* = 0.038, best modulation frequency: *p* = 0.732).

**Figure 5:**
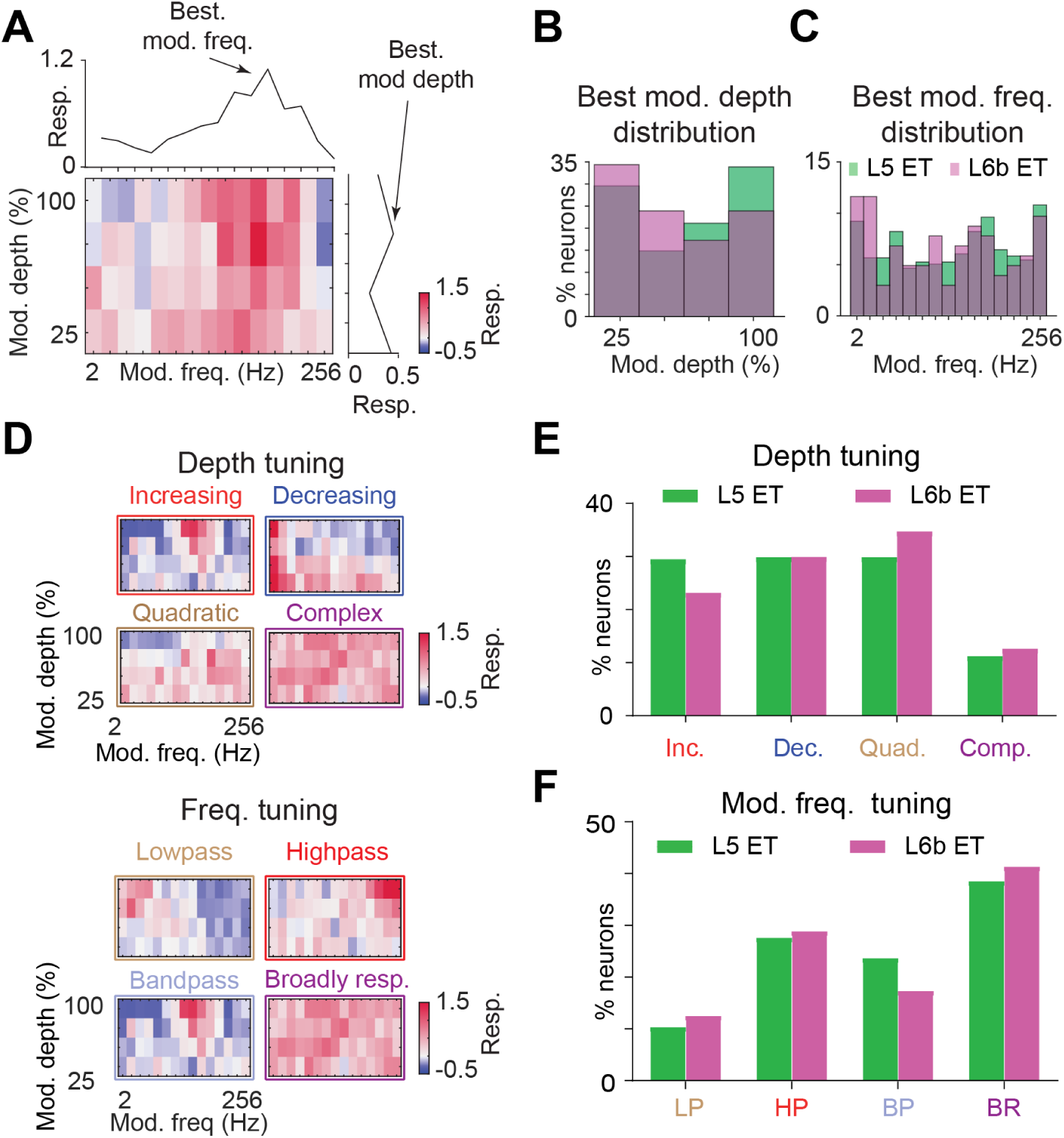
L5 and L6b ET neurons show similar sAM noise tuning profiles. **A**: Example modulation transfer function (MTF) with modulation frequency on the x-axis and modulation depth on the y-axis. Top, depth-averaged response used to identify best modulation frequency (peak). Right, frequency-averaged response used to identify best modulation depth (peak). **B-C**: Distribution of best modulation depth **(B)** and best modulation frequency **(C)** for L5 and L6b ET neurons. **D**: Top, example MTFs depicting increasing (upper left), decreasing (upper right), quadratic (lower left), and complex (lower right) modulation depth tuning profiles. Bottom, example MTFs depicting lowpass (upper left), highpass (upper right), bandpass (lower left), and broadly responsive (lower right) modulation frequency tuning profiles. **E-F**: Distribution of modulation depth **(E)** and modulation frequency **(F)** tuning categories for L5 and L6b ET neurons. No significant differences were observed between subtypes (chi-squared test, see Results for statistics).

As in Figure 4, we clustered each neuron’s modulation depth and frequency tuning curves. The depth tuning profiles were classified into four categories: increasing, decreasing, quadratic (U-shaped), and complex (**Figure 5D** *top*). Modulation frequency tuning curves were classified as: lowpass, highpass, bandpass, or broadly responsive (**Figure 5D** *bottom*). Both L5 ET and L6b ET neurons exhibited a diverse mix of depth and frequency tuning patterns. However, the proportion of L5 ET and L6b ET neurons in each tuning category did not differ significantly between subtypes (**Figure 5E-F**, chi-squared test, depth tuning: *p* = 0.174, frequency tuning: *p* = 0.797). Thus, unlike the clear subtype-specific differences observed for pure tones, L5 ET and L6b ET neurons showed comparable tuning to sAM noise, suggesting that temporal modulation processing is a shared feature of both subpopulations.

### L5 and L6b ET neurons exhibit similar spectrotemporal modulation tuning to ripples

To probe sensitivity to joint spectrotemporal modulation beyond simple tonal and envelope features, we presented ripple stimuli that systematically varied spectral density and temporal modulation rate (**Figure 1H**) [41, 42]. Neuronal responses to these stimuli were used to compute a ripple transfer function (RTF) for each sound-excited neuron, which quantified sensitivity to joint spectrotemporal modulations [42]. From each RTF, we calculated the best spectral and temporal modulation values, defined as the stimulus parameters that evoked the strongest responses averaged across temporal or spectral modulation, respectively (**Figure 6A**).

**Figure 6:**
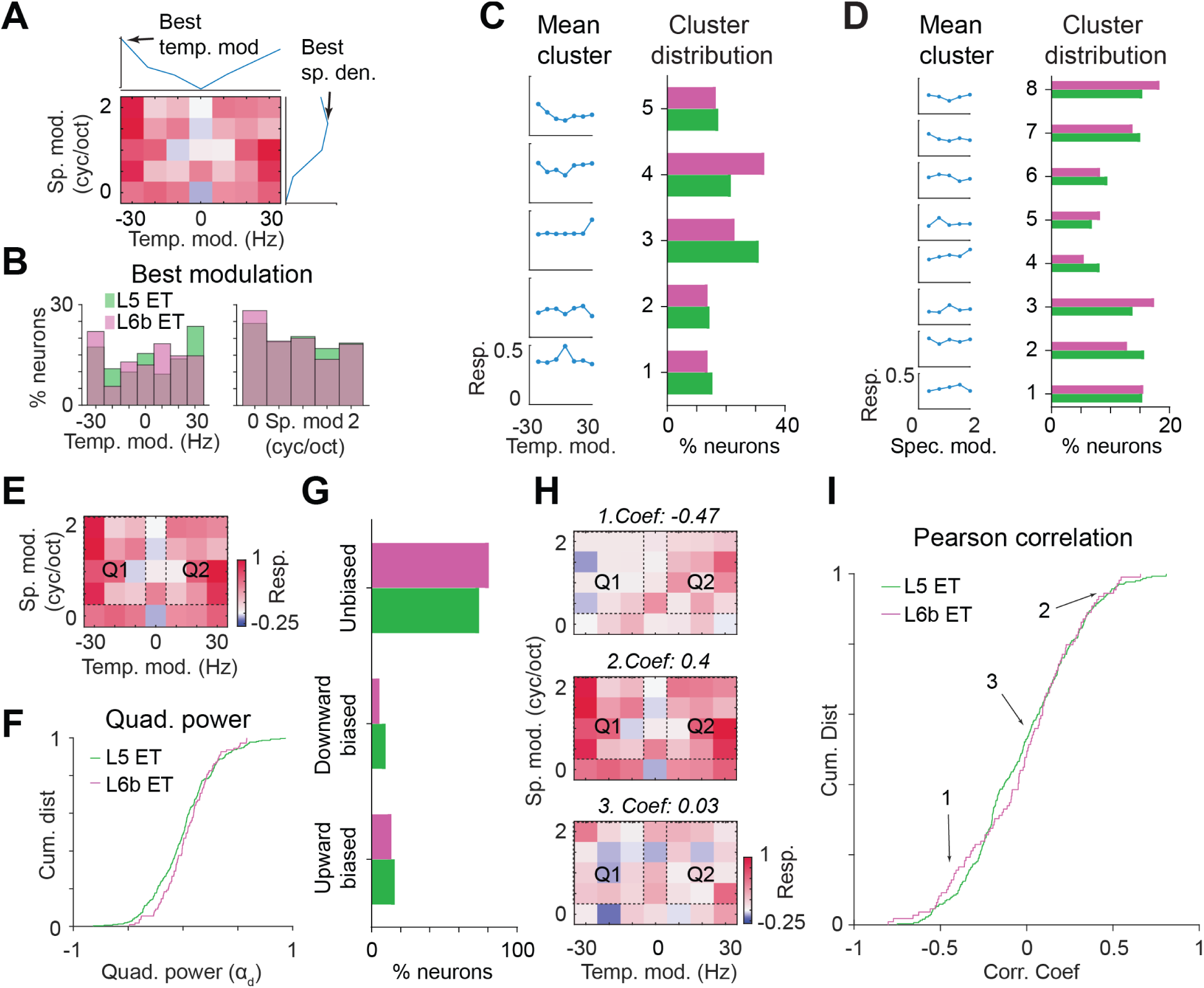
Spectrotemporal ripple tuning is largely similar between L5 and L6b ET neurons. **A**: Example ripple transfer function (RTF) with temporal modulation rate on the x-axis and spectral modulation density on the y-axis. Top, spectral-modulation-averaged response used to identify best temporal modulation (peak). Right, temporal-modulation-averaged response used to identify best spectral modulation (peak). **B**: Distribution of best temporal modulation (Hz) and best spectral modulation (cycles/octave) for L5 and L6b ET neurons. **C**: Mean temporal modulation tuning profiles for each cluster (left) and proportion of L5 and L6b ET neurons in each cluster (right). **D**: Mean spectral modulation tuning profiles for each cluster (left) and proportion of L5 and L6b ET neurons in each cluster (right). **E**: Example RTF from a ripple-responsive neuron illustrating the two quadrants (Q1 and Q2) corresponding to opposite temporal modulation directions, used for the quadrant power analysis (see Methods). **F**: Distribution of quadrant power for L5 and L6b ET neurons. **G**: Distribution of quadrant biases for L5 and L6b ET neurons. **H**: Example RTFs with negative (top), positive (middle), and near-zero (bottom) Pearson correlations between Q1 and Q2 responses. **I**: Distribution of Pearson correlation coefficients between Q1 and Q2 responses for L5 and L6b ET neurons. Numbered outlines correspond to the examples in panel **H**. No significant differences were observed between subtypes (Kolmogorov-Smirnov test, see Results for statistics).

L5 and L6b ET neurons responded similarly to ripple stimuli but differed in their temporal modulation preferences. L5 ET neurons preferred fast-moving upward and downward moving ripples, whereas L6b ET neurons showed a bias toward fast downward and slower upward moving ripples (**Figure 6B**, chi-squared test, *p* = 0.034). In contrast, the distribution of spectral modulation preferences did not differ between the two subtypes (**Figure 6B**, chi-squared test, *p* = 0.907). Together, these results indicate that subtype differences are evident in preferred temporal modulation parameters, whereas spectral modulation preferences are similar. We next asked whether these differences extend to broader tuning-curve motifs beyond the preferred parameter values. We therefore performed unsupervised clustering of temporal and spectral tuning profiles as in Figures 4 and 5. This analysis revealed 5–8 distinct tuning motifs in each domain, but their distribution did not differ significantly between L5 and L6b ET neurons (**Figure 6C-D**, chi-squared test, temporal modulation: *p* = 0.177, spectral modulation: *p* = 0.900).

Given subtype differences in preferred temporal modulation parameters (**Figure 6B**), we next asked whether L5 and L6b ET neurons differ in directional selectivity for upward-versus downward-drifting ripples. We quantified direction selectivity using a quadrant power analysis [42], where positive values indicated a bias for upward sweeps, negative values for downward sweeps, and values near zero indicated overall symmetric responses (**Figure 6F-G**). L5 and L6b ET neurons did not show significant differences in the distribution of quadrant power, suggesting that both populations respond similarly to upward- and downward-drifting ripple stimuli (**Figure 6F-G**, Kolmogorov-Smirnov test, *p* = 0.144). As a complementary measure of directional symmetry, we calculated the Pearson correlation between the neural responses to the matched upward-versus downward-drifting stimuli in quadrants 1 and 2. We found that the distribution of correlation coefficients was broadly similar between L5 and L6b ET subtypes (**Figure 6H-I**, Kolmogorov-Smirnov test, *p* = 0.324). Together, these findings show that L5 and L6b ET neurons exhibit similar directional symmetry in ripple-evoked responses, despite subtype-specific differences in preferred temporal modulation parameters.

### L5 ET neurons exhibit higher pure-tone response sparseness than L6b ET neurons

L5 and L6b ET neurons differed most prominently in their responses to pure tones. Whereas most L5 ET neurons had monotonic and single-peaked FRAs, L6b ET neurons showed a greater prevalence of non-monotonic and complex FRAs. These qualitative differences suggest that L5 responses are restricted to fewer pure-tone stimuli across frequency and sound level, whereas L6b responses are distributed across more of the stimulus ensemble. We therefore quantified response sparseness, a standard measure of how a neuron’s responses are distributed across stimuli (i.e., the extent to which it responds to many stimuli versus a small subset) [8, 43, 44]. The sparseness index ranged from 0 to 1, where values close to 0 indicate responses distributed broadly across stimuli, and values near 1 reflect highly selective responses (**Figure 7A**). Because ET neurons exhibited both sound-evoked increases and decreases in activity, we included both sound-excited and sound-suppressed neurons in the sparseness analysis. L5 ET neurons exhibited significantly higher sparseness indices than L6b ET neurons, indicating that they respond to a narrower subset of pure tone stimuli (**Figure 7B**, Wilcoxon rank-sum test, *p* = 4.7 × 10^−13^). This difference was driven entirely by sound-excited neurons (**Figure 7B**, Wilcoxon rank-sum test, *p* = 0.002). In contrast, sound-suppressed neurons from both L5 and L6b responded to a broad range of stimuli, with sparseness indices close to 0 (**Figure 7B**, Wilcoxon rank-sum test, *p* = 0.537). These results suggest that L5 ET neurons are more selective in their frequency-intensity tuning, consistent with their predominantly single-peaked receptive fields, whereas L6b ET neurons show lower selectivity among sound-excited responses, consistent with their more complex FRAs.

**Figure 7:**
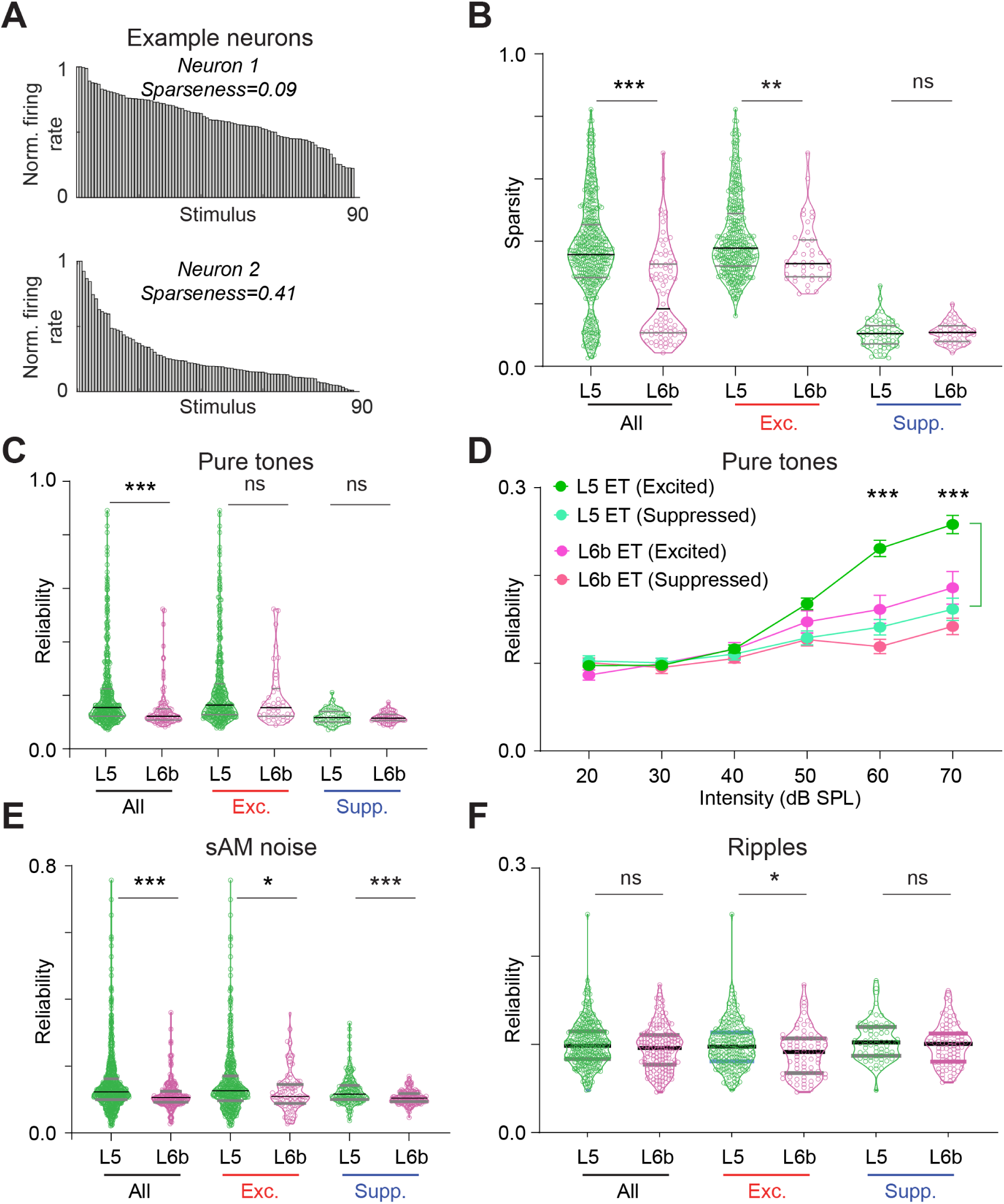
Response sparseness and reliability are higher in L5 than L6b ET neurons. **A**: Example normalized response distributions illustrating low (top) and high (bottom) sparseness indices. **B**: Sparseness indices for L5 and L6b ET neurons, shown separately for all sound-responsive, sound-excited, and sound-suppressed neurons. **C**: Reliability indices for all sound-responsive, sound-excited, and sound-suppressed ET neurons in response to pure tones. **D**: Reliability as a function of sound intensity for sound-excited (left) and sound-suppressed (right) L5 and L6b ET neurons in response to pure tones. **E**: Reliability indices in response to sAM noise, shown separately for all sound-responsive, sound-excited, and sound-suppressed subgroups. **F**: Same as E for spectrotem-poral ripples. Asterisks denote statistically significant differences using Wilcoxon rank-sum test **(B, C, E, F)** and Tukey’s post-hoc test after two-way ANOVA **(D)**. * *p* ≤ 0.05, ** *p* ≤ 0.01, *** *p* ≤ 0.001, ns = non-significant.

### L5 ET neurons exhibit higher response reliability than L6b ET neurons

ACtx neurons vary in their responses to repeated presentations of the same sound, influenced by internal state, synaptic dynamics, and adaptation [45–48]. To quantify the trial-to-trial variability of ET responses, we computed a reliability index for each sound-responsive neuron. This index ranged from 0 to 1, with values near 1 indicating highly reliable responses and values near 0 reflecting high variability. Pure-tone responses in L5 ET neurons were more reliable than those in L6b ET neurons (**Figure 7C**, Wilcoxon rank-sum test, *p* = 1.0 × 10^−7^). To assess intensity dependence, we calculated response reliability at each presented sound level. In sound-excited neurons, reliability increased with rising intensity in L5 ET neurons, and subtype-specific differences emerged at higher intensities (**Figure 7D**, two-way ANOVA, main effect of intensity and subtype: *p* = 1.0 × 10^−15^ and 8.3 × 10^−14^, respectively, intensity × subtype interaction: *p* = 2.6 × 10^−10^, Tukey’s post-hoc test, *p* = 3.5 × 10^−4^ at 60 and 1.9 × 10^−4^ at 70 dB SPL), whereas differences were non-significant at lower intensities (**Figure 7D**, Tukey’s post-hoc test, 20–50 dB SPL, *p >* 0.05 for each intensity-matched pair). Among pure-tone suppressed neurons, reliability showed only a weak dependence on intensity, and no significant differences were observed between L5 and L6b subtypes (**Figure 7D**, Tukey’s post-hoc test, *p >* 0.05 for each intensity-matched pair).

Likewise, L5 ET responses to sAM noise were more reliable than L6b ET responses (**Figure 7E**, Wilcoxon rank-sum test, *p* = 9.02 × 10^−9^). When reliability indices were stratified by response type, both sound-excited and sound-suppressed L5 ET neurons showed significantly higher reliability than L6b ET neurons (**Figure 7E**, Wilcoxon rank-sum test, sAM excited, *p* = 0.020, sAM suppressed, *p* = 2.3 × 10^−6^). In contrast, responses to ripples were uniformly less reliable across both subtypes, and no significant difference between subtypes was observed across sound-responsive groups (**Figure 7F**, Wilcoxon rank-sum test, *p >* 0.05 for all sound-responsive groups). Collectively, these results indicate that L5 ET neurons exhibit greater trial-to-trial reliability than L6b ET neurons, particularly for pure tones and sAM noise, supporting more consistent stimulus-evoked responses. Lower reliability in L6b ET neurons may reflect greater variability associated with network state or other contextual influences.

### L6b ET neurons exhibit stronger functional coupling than L5 ET neurons

Shared trial-to-trial variability between neurons, commonly measured via noise correlations, is thought to reflect underlying functional connectivity and can constrain the amount of independent information a population can encode [49–51]. We computed pairwise noise correlations to quantify shared variability within L5 and L6b neuronal responses to auditory stimuli (**Figure 8A-B**). Correlation coefficients were calculated between neuron pairs within each field-of-view, stratified as sound-excited pairs, sound-suppressed pairs, all responsive pairs (excited or suppressed), or all pairs regardless of response type. Within L5 ET neurons, pairwise noise correlations were uniformly low, with no significant difference between excited and suppressed neurons (**Figure 8C**, Kolmogorov-Smirnov test, *p* = 0.076). In contrast, within L6b ET neurons, noise correlations among sound-suppressed pairs were significantly higher than among sound-excited pairs (**Figure 8D**, Kolmogorov-Smirnov test, *p* = 2.7 × 10^−64^).

**Figure 8:**
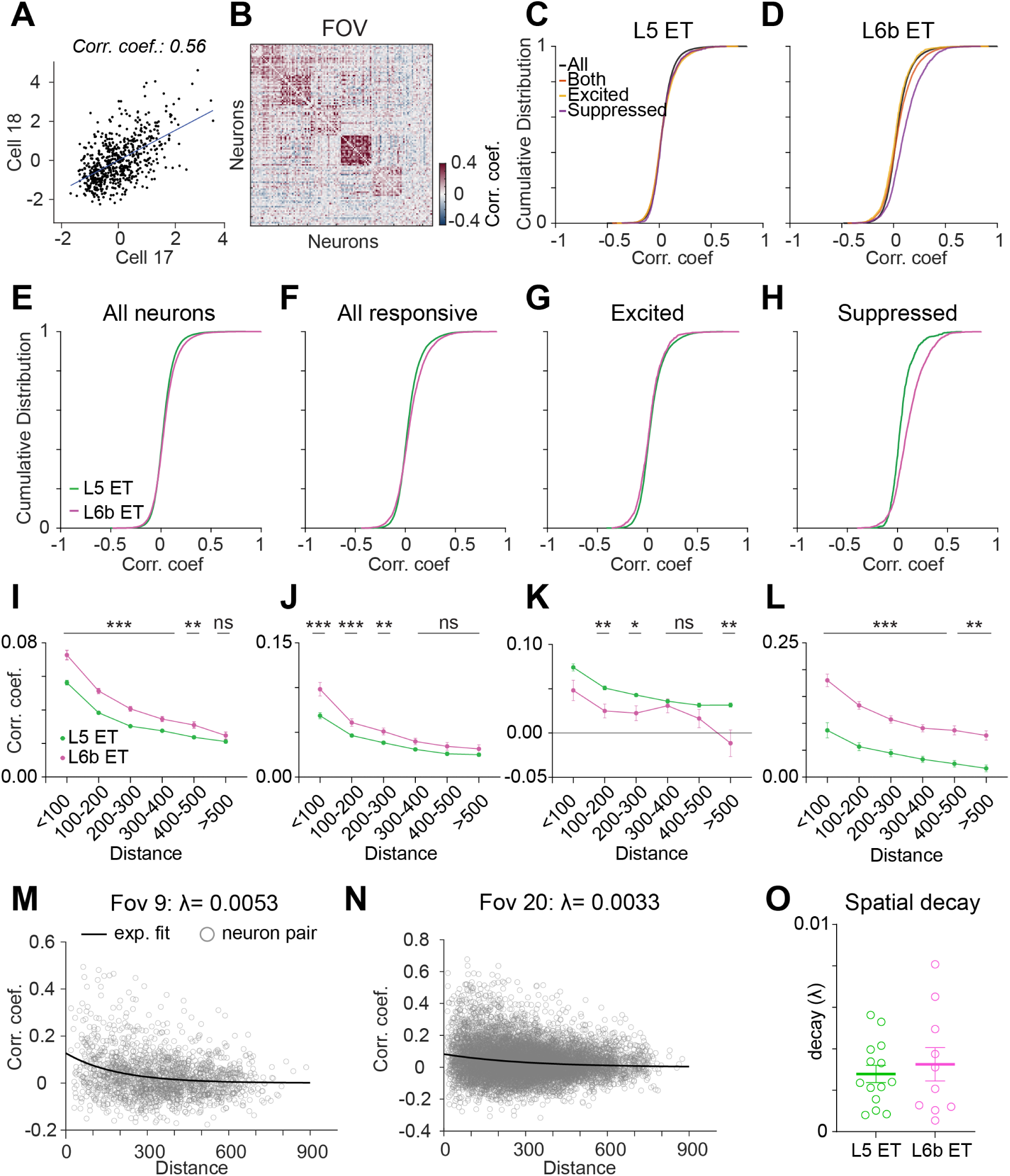
L6b ET neurons show stronger pairwise noise correlations than L5 ET neurons. **A**: Example pairwise noise correlation between a pair of L5 ET neurons. **B**: Example pairwise noise correlation matrix for all simultaneously recorded neurons within a single FOV. Larger correlation coefficients are organized along the diagonal for visualization. **C-D**: Distribution of pairwise correlation coefficients for all recorded, all sound-responsive, sound-excited, and sound-suppressed neuron pairs within L5 **(C)** and L6b **(D)**. **E-H**: Comparison of correlation coefficient distributions between L5 and L6b ET neurons for all recorded pairs **(E)**, all sound-responsive pairs **(F)**, sound-excited pairs **(G)**, and sound-suppressed pairs **(H)**. **I-L**: Pairwise noise correlations as a function of intersomatic distance for all recorded **(I)**, all sound-responsive **(J)**, sound-excited **(K)**, and sound-suppressed **(L)** L5 and L6b ET neuron pairs. **M-N**: Example FOVs illustrating exponential decay fits with higher **(M)** and lower **(N)** spatial decay constants. **O**: Comparison of spatial decay constants between L5 and L6b ET neurons. Asterisks denote statistically significant differences using Šidák’s post-hoc test after two-way ANOVA. * *p* ≤ 0.05, ** *p* ≤ 0.01, *** *p* ≤ 0.001, ns = non-significant.

We next compared noise correlation magnitudes between L5 and L6b populations. Overall, L6b ET neurons exhibited significantly stronger correlated activity than L5 ET neurons, both when considering all neurons (**Figure 8E-F**, Kolmogorov-Smirnov test, *p* = 5.8×10^−51^) and when restricting the analysis to responsive neurons only (**Figure 8E-F**, Kolmogorov-Smirnov test, *p* = 3.3 × 10^−28^). Interestingly, the small fraction of L6b ET neurons that were sound-excited showed especially low noise correlations, lower than those of L5 sound-excited pairs (**Figure 8G**, Kolmogorov-Smirnov test, *p* = 2.7 × 10^−6^). Conversely, sound-suppressed L6b ET neurons were more strongly correlated than their L5 counterparts (**Figure 8H**, Kolmogorov-Smirnov test, *p* = 8.4 × 10^−53^). These findings suggest that suppressed L6b ET neurons form a more tightly coupled functional network than L5 ET neurons, while sound-excited L6b ET neurons respond more independently.

Previous studies in sensory cortex have shown that pairwise correlations generally decrease with increasing intersomatic distance; however, the strength of spatial coupling may vary by neuron type [52–54]. To assess spatial coupling differences between ET subtypes, we binned neuron pairs by intersomatic distance. L6b ET neurons showed stronger noise correlations than L5 ET pairs at most intersomatic distance bins (**Figure 8I**, two-way ANOVA, main effect of subtype: *p* = 1.0 × 10^−15^, main effect of distance: *p* = 1.0 × 10^−15^, subtype × distance interaction: *p* = 3.0 × 10^−4^). This pattern held when restricting to all sound-responsive pairs (**Figure 8J**, two-way ANOVA, main effect of subtype: *p* = 5.6 × 10^−12^, main effect of distance: *p* = 1.0 × 10^−15^, subtype × distance interaction: *p* = 0.039), and for sound-excited pairs alone (**Figure 8K**, two-way ANOVA, main effect of subtype: *p* = 7.7 × 10^−10^, main effect of distance: *p* = 2.7 × 10^−9^, subtype × distance interaction: *p* = 0.091). The difference was largest and most consistent among sound-suppressed pairs (**Figure 8L**, two-way ANOVA, main effect of subtype: *p* = 1.0×10^−15^, main effect of distance: *p* = 1.0 × 10^−15^, subtype × distance interaction: *p* = 0.451). These differences were especially pronounced at short intersomatic distances (**Figure 8I**, all neuron pairs, two-way ANOVA with Šidák’s post-hoc test, 0–100 *µ*m: *p* = 2.1 × 10^−12^, 100–200 *µ*m: *p* = 1.0 × 10^−15^, 200–300 *µ*m: *p* = 1.8 × 10^−12^), consistent with our earlier observation of elevated shared variability among L6b ET neurons, particularly in the sound-suppressed population (**Figure 8H**). To quantify the spatial extent of correlated variability, we fitted exponential decay functions and extracted a decay constant for each subtype (**Figure 8M-N**). Decay constants did not differ between L5 and L6b ET neurons (**Figure 8O**, Wilcoxon rank-sum test, *p* = 0.977). Overall, L6b ET neurons exhibited higher noise correlations than L5 ET neurons at most intersomatic distances tested, with both populations showing a similar rate of distance-dependent decay.

## Discussion

In this study, we characterized the *in vivo* sensory response properties of ACtx L5 and L6b ET neurons across a range of acoustic stimuli. We found that L5 ET neurons were more frequently excited by sound, showed higher trial-to-trial response reliability, and exhibited sharper tuning to pure tones. In contrast, L6b ET neurons were more often suppressed by sound, demonstrated more complex and broader tuning profiles, and displayed stronger pairwise correlated activity. These subtype-specific differences, spanning response sign, selectivity, reliability, and functional coupling (noise correlations), are reminiscent of parallel processing architectures in other sensory systems [55–57] and suggest that L5 and L6b ET neurons support complementary corticofugal channels: one conveying sparse, reliable representations of salient acoustic features (L5), and the other providing broader, more integrative signals that may be shaped by behavioral and neuromodulatory state (L6b).

### Distinct excitatory and suppressive response profiles of L5 and L6b ET neurons

L5 ET neurons were predominantly excited by sound, with a substantial fraction showing robust excitation across all stimulus types (**Figure 2E**). In contrast, L6b ET neurons showed a more balanced mix of excitation and suppression, with a markedly higher proportion displaying suppressive responses, particularly to complex stimuli such as sAM noise and spectrotemporal ripples. This subtype-specific divergence in response profiles likely arises from differences in anatomical connectivity and input sources. L5 ET neurons, with prominent apical dendrites extending into superficial layers, are well-positioned to integrate direct thalamocortical and intracortical inputs, along with higher-order feedback. L6b ET neurons, however, are morphologically diverse, including pyramidal-like and oval-shaped cells with radially oriented somata and dendrites that branch profusely through deeper layers [10, 12], where they likely receive predominantly intracortical input. Indeed, L6 corticofugal neurons receive prominent feedforward inhibitory inputs that dynamically shape their responses, often resulting in sound-evoked suppression [25, 58]. Although these prior observations derive from studies of L6a corticothalamic neurons rather than L6b ET neurons, prominent inhibitory drive may be a shared feature of deep L6 corticofugal circuits more broadly.

Beyond these differences in overall excitation and suppression, both L5 and L6b ET neurons displayed a range of temporal response motifs, including onset, sustained, and offset patterns, reflecting diverse temporal processing within ACtx (**Figure 3B-E**). Onset responses were more common in L5 ET neurons, consistent with a role in alerting downstream targets such as the IC or SC to the presence of new sounds [59, 60]. Sustained responses, also prevalent in L5, may encode stimulus duration or support perceptual continuity and experience-dependent auditory learning [61–63]. In comparison, many L6b ET neurons displayed suppression lasting at least the duration of the stimulus, or complex multiphasic responses to modulated stimuli. Such suppression may arise from reversed synaptic integration, in which strong inhibitory inputs precede excitatory inputs, effectively silencing L6 corticofugal neurons [25]. How these distinct temporal profiles influence processing in downstream targets remains an open question. The net effect on subcortical structures such as the IC will depend on the identity and circuit role of the postsynaptic neurons contacted by each ET subtype. Given the considerable cell-type diversity within the IC, resolving this will require circuit-level investigations that map the synaptic targets of L5 and L6b ET projections.

### Subtype-specific tuning to pure tones but not complex sounds

Among sound-excited neurons, the most prominent subtype-specific differences emerged in responses to pure tones. L5 ET neurons predominantly exhibited monotonic intensity tuning and single-peaked frequency tuning profiles. In contrast, L6b ET neurons showed non-monotonic intensity tuning and complex, multi-peaked frequency tuning (**Figure 4F-G**). These broader receptive fields in L6b ET neurons could reflect integration across a wider range of intracortical inputs. Indeed, L6b neurons receive long-range excitatory and inhibitory connections from secondary auditory areas and other sensory regions, which could contribute to tuning properties that diverge from the tonotopic organization typical of primary ACtx [64, 65].

In contrast to the clear subtype-specific differences observed for pure tones, L5 and L6b ET neurons exhibited largely similar tuning profiles for more complex sounds, including sAM noise and spectrotemporal ripples (**Figures 5E-F, 6C-D**). One possibility is that temporal and spectrotemporal modulation tuning relies on circuit mechanisms that are shared across cortical layers, such as synaptic depression or inhibitory network dynamics, and are therefore less sensitive to the laminar differences in input that shape pure-tone selectivity. This convergence may also reflect the broadband nature of these stimuli: unlike pure tones, which probe frequency-specific channels where L5 and L6b inputs diverge, sAM noise and ripples activate neurons across many frequency channels simultaneously, potentially masking the selectivity differences that distinguish the two subtypes. Moreover, temporal modulation features may be substantially shaped by subcortical processing prior to reaching cortex, such that both ET subtypes inherit similar modulation representations from shared thalamocortical inputs. Collectively, these results suggest that L5 and L6b ET neurons participate in functionally distinct corticofugal pathways for simple sounds: L5 ET neurons convey precise frequency-intensity representations, whereas L6b ET neurons convey broader, multi-frequency representations that may reflect convergent intracortical integration. However, these coding differences appear to be most pronounced for pure tones and may not extend to more complex acoustic features.

Calcium imaging as used in this study does not capture subthreshold integration, burst firing patterns, or fine temporal dynamics due to its limited temporal resolution, potentially masking differences in spike timing or rapid response dynamics [66, 67]. This limitation is particularly relevant for stimuli that engage temporal processing, such as sAM noise and spectrotemporal ripples. The largely similar tuning profiles of L5 and L6b ET neurons for these complex sounds may therefore partly reflect this methodological constraint rather than a true absence of subtype-specific coding differences. Future work using targeted electrophysiological recordings could determine whether L5 and L6b ET neurons exhibit distinct firing patterns for these stimuli, as differences in action potential kinetics and bursting behavior have been noted between L5 ET and other deep-layer neuronal subclasses *in vitro* [12].

### Divergent coding strategies: sparseness and reliability

L5 ET neurons encoded pure tones with higher sparseness than L6b ET neurons (**Figure 7B**). Sparse coding, in which neurons respond selectively to a small subset of stimuli, is generally considered an efficient strategy for transmitting information with minimal redundancy [43, 68]. In contrast, the lower sparseness of L6b ET neurons, reflecting broader responses across the stimulus ensemble, is consistent with integration across a wider range of intracortical inputs from diverse tonotopic channels. This distinction parallels the tuning differences described above (**Figure 4**) and reinforces the view that L5 and L6b ET neurons occupy different positions along a selectivity-integration axis. Prior studies of L6 corticothalamic neurons have found that they exhibit sharper pure-tone tuning and suppress the activity of nearby neurons in the cortical column [8, 69, 70]. However, L6b ET neurons belong to a distinct circuit that projects to higher-order thalamus and other subcortical targets [10]. The broader tuning and lower sparseness we observe in L6b ET neurons therefore cannot be predicted from classical L6 corticothalamic function, and likely reflect the distinct input-output architecture of this population.

Trial-to-trial reliability of neural responses to pure tones and sAM noise also differed between ET subtypes (**Figure 7C-F**). Reliability was particularly high in L5 ET neurons at higher sound intensities, consistent with a role in conveying robust auditory signals to downstream targets during sensory-guided behaviors [71, 72]. In contrast, L6b ET neurons exhibited lower reliability across all stimulus classes tested. Lower reliability can arise from synaptic noise, intrinsic neuronal properties such as membrane potential fluctuations [45, 73], and external influences including attention, arousal, and ongoing network dynamics [47, 48, 74]. The lower reliability of L6b ET neurons may therefore reflect a greater sensitivity to brain state [29], raising the possibility that these neurons require specific neuromodulatory or task-dependent facilitation to respond consistently. Our passive listening paradigm did not engage goal-directed behavior, and it is possible that L6b ET response reliability would increase under conditions that recruit top-down or neuromodulatory inputs to deep cortical layers.

### Stronger functional coupling among L6b ET neurons

Noise correlations were significantly higher among L6b ET neurons than L5 ET neurons, particularly among sound-suppressed pairs (**Figure 8E-H**). This elevated shared variability suggests that L6b ET neurons are more tightly functionally coupled, potentially reflecting shared input sources or recurrent connectivity. High noise correlations can reduce the independent information carried by individual neurons, limiting the details of acoustic feature representations conveyed to subcortical targets [49, 75], yet may support population-level operations such as gain modulation or coordinated state transitions [76, 77]. The observation that suppressed L6b ET neuron pairs showed the highest correlations, while the small fraction of excited L6b pairs showed especially low correlations (lower even than L5 excited pairs), suggests that these two response classes within L6b may constitute functionally distinct subpopulations with different network integration profiles.

One potential source of elevated correlations among L6b ET neurons is convergent input from higher-order cortical areas or neuromodulatory centers that synchronize activity across the population. Indeed, L6b neurons are highly sensitive to neuromodulators such as acetylcholine, dopamine, and orexin, which regulate brain states such as arousal and alertness [22, 29]. Elevated noise correlations in the passive, awake state may therefore reflect tonic neuromodulatory drive that acts broadly across the L6b population. Whether these correlations decrease under conditions of active engagement, when neuromodulatory inputs may shift from tonic to phasic modes, remains an open question. In contrast, the lower noise correlations among L5 ET neurons indicate that individual neurons carry more independent information [49, 75], enabling the population to transmit a richer, more diverse set of acoustic feature representations to subcortical targets.

Collectively, these findings indicate that L5 and L6b ET neurons form complementary corticofugal processing streams from ACtx to subcortical targets [1, 8–10]. L5 ET neurons convey sparse, selective, and reliable representations of acoustic features, particularly pure tones. In contrast, L6b ET neurons exhibit broader, less selective tuning, more prominent sound-evoked suppression, lower trial-to-trial reliability, and stronger functional coupling within the local population. These properties are consistent with a corticofugal channel whose output is shaped less by individual stimulus identity and more by network state and convergent intracortical input [76, 77]. Together, these parallel output streams may enable the ACtx to simultaneously provide subcortical targets with both precise acoustic feature information and broader, state-dependent contextual signals, thereby enriching the repertoire of descending control over subcortical auditory processing.

## Materials and Methods

### Mice

All procedures were approved by the University of Pittsburgh Animal Care and Use Committee (Protocol #: 25036474) and followed the guidelines established by the National Institutes of Health for the care and use of laboratory animals. We used a total of 10 adult wild-type C57BL/6J mice (#000664, Jackson Labs) of either sex in this study. All mice were light-reversed (12 h dark/light cycle) with ad libitum access to food and water throughout experiments. All imaging was conducted during the dark cycle.

### Surgical Procedures

All surgical procedures were conducted in anesthetized mice under aseptic conditions in a stereotaxic frame (Kopf model 1900). Mice were anesthetized with 5% isoflurane in oxygen and maintained at 1-2% throughout surgery. Mice lay atop a homeothermic blanket system (Fine Science Tools) that maintained core body temperature at approximately 36.5^◦^C. Lidocaine hydrochloride (2%, local anesthetic) was injected subcutaneously prior to skin incision. Ophthalmic ointment was applied over the eyes to prevent dryness. After surgery, mice received a subcutaneous injection of an analgesic (carprofen, 5 mg/kg) and a carprofen-infused MediGel (Clear H_2_O) was provided in their home cages for three days post-operation. All mice underwent two surgical procedures: viral delivery and cranial window implantation.

#### Virus-mediated gene delivery

All virus injections were conducted on mice aged 6-10 weeks. An approximately 1-cm-long midline incision was made to expose the skull. A ∼0.2-0.3 mm diameter burr hole was drilled over the right hemisphere at AP −5 mm, ML −0.9 mm to target the right IC. A motorized stereotaxic injection system (Nanoject III, Drummond Scientific) was used to deliver a total of 500 nl (250 nl at each depth) of CAV-2 Cre (Plateforme de Vectorologie de Montpellier, titer: 1×10^12^ vg/ml) at depths of 450 and 900 *µ*m below the pial surface. For viral delivery to ACtx, incisions were made on the right side of the scalp to expose the skull around the caudal end of the temporal ridge. The temporalis muscle was retracted and two burr holes (∼ 0.3 mm in diameter) were drilled along the temporal ridge, spanning 1-2 mm rostral to the lambdoid suture. 250 nl of Cre-dependent GCaMP8s (pGP-AAV1-syn-FLEX-jGCaMP8s-WPRE, Addgene #162377-AAV1, titer: 7 × 10^12^ vg/ml) was injected at a depth of 550 µm below the pial surface in each burr hole. Viruses were diluted with PBS to achieve the desired titer and the rate of injection was 10 nl every 45-60 seconds. The glass pipette remained in the brain for an additional 10 minutes at each site following virus delivery for all injections. Incision sites were sutured and antibiotic ointment was applied.

#### Cranial window implantation

Three weeks following virus injection, mice were anesthetized for chronic imaging window implantation surgery. An intraperitoneal injection of dexamethasone sodium phosphate (2 mg/kg) was administered to reduce inflammation and brain swelling. The scalp overlying the dorsal skull was removed and an etchant (C&B Metabond) was applied to the dorsal surface of the skull to facilitate head plate adhesion. A customized titanium head plate (eMachineShop) was then affixed to the skull using dental cement (C&B Metabond). A 3-mm-diameter craniotomy was made over the right ACtx. The overlying dura was carefully removed and the exposed brain was flushed with cold saline. A cranial window, comprising a stack of three glass coverslips (two 3-mm and one 4-mm diameter, bonded with UV-curing optical adhesive (#68, Norland), was then inserted such that the 3-mm coverslips fit within the craniotomy and the 4-mm coverslip rested on the surrounding skull. The gap between the skull and window was sealed with silicone elastomer (Kwik-Sil, World Precision Instruments) and the window was secured to the skull using dental cement. The remaining skin edges were attached to the hardened cement using tissue adhesive (Vetbond, 3M).

### Acoustic stimulation

Sounds were generated with a 24-bit digital-to-analog converter (PXI4461, National Instruments) using custom scripts in MATLAB (MathWorks) and LabVIEW (National Instruments). Acoustic stimuli were delivered using a free-field speaker (PUI Audio), ∼20 cm from the left (contralateral) ear and calibrated using a free-field prepolarized microphone (377C01, PCB Piezotronics). Three types of acoustic stimuli were presented, each in pseudorandom order, with 2-2.5 sec inter-stimulus intervals and 10-15 repetitions per unique stimulus condition. Pure tones (50 or 500 ms duration, with 5 ms cosine-squared onset/offset ramps) ranged from 4 to 45 kHz in 0.25-octave steps at intensities between 20 and 70 dB SPL. sAM noise (500 ms, at 70 dB SPL) was presented at modulation frequencies from 2 to 256 Hz (0.5 octave spacing) and modulation depths of 25%, 50%, 75%, and 100%. Spectrotemporal ripples (500 ms, spanning 4 to 45 kHz at 70 dB SPL) varied in spectral modulation density from 0 to 2 cycles per octave (0.5 cyc/oct spacing) and temporal modulation rate from −30 to +30 Hz (10 Hz spacing, where negative and positive values indicate downward- and upward-drifting spectral envelopes, respectively).

### Two-photon calcium imaging

One week after cranial window implantation, mice were acclimated to the head-fixed assembly for 2-3 days. Neural activity in response to four pure tones (4, 8, 16, and 32 kHz) was captured by widefield fluorescence imaging (Bergamo, ThorLabs) and used to functionally confirm the location of the right primary ACtx [78]. The two-photon (2P) field of view was centered over primary ACtx and imaging of L5 and L6b ET neurons was performed in passively listening mice. All imaging was conducted within the same time window of the animals’ light cycle to minimize circadian variability. Two-photon calcium imaging was conducted using an InSightX3 (Spectra Physics) laser tuned to 940 nm and a water-immersion objective (Nikon 16x). All two-photon imaging (Bergamo, ThorLabs) was of the right ACtx. Mice were head-fixed upright with the microscope rotated to be parallel to the cranial window (approximately 40 to 60^◦^ tilt). Images were collected at 30 Hz. Imaging was performed in a dark, sound-attenuating chamber, and the mice were monitored during experiments using an infrared camera (Genie Nano, Teledyne). L5 and L6b ET neurons were imaged at 400-550 and 700-830 *µ*m below the pial surface, at laser powers of 55-120 and 100-150 mW, respectively. Fluorescence images were captured at either 1x (749 × 749 *µ*m) or 2x (374.5 × 374.5 *µ*m) digital zoom.

### Quantification and data analysis

#### Two-photon

Post-recording analysis began with Suite2p [**pachitariu^·^2016**], an open-source image processing pipeline. Suite2p was used to register raw calcium movies, detect spatial regions of interest (ROIs), classify neuronal and non-neuronal ROIs, extract calcium fluorescence signals, and perform spike deconvolution. All ROIs were then manually inspected and only those corresponding to neuronal somata were retained for analysis. Deconvolved spike estimates were z-scored by subtracting the mean and dividing by the standard deviation (SD) of activity during a 500 ms pre-stimulus baseline window.

#### Responsiveness, receptive fields, and response tuning

Neurons were classified as sound-excited if their deconvolved spike z-score exceeded 2 SD above mean baseline activity (calculated from the 500-ms pre-stimulus window) in at least 3 consecutive frames within the response window (15–30 frames (500–1000 ms) for 50 ms pure tones and 15–40 frames (500–1333 ms) for 500 ms sounds) on at least 50% of trials for the stimulus evoking the strongest response (best stimulus). Sound-evoked suppression was typically weaker in magnitude than excitation, and the stimulus evoking maximal suppression was less clearly defined. Given the smaller magnitude of suppressive responses in calcium imaging, a lower threshold was used: neurons were classified as sound-suppressed if, in response to the stimulus that evoked the strongest suppression, their activity dropped by at least 1 SD below baseline in at least 3 consecutive frames within the response window on more than 50% of trials (adapted from [79]). Receptive fields were constructed for each neuron and stimulus class. The mean response to each stimulus was computed within a 10-frame window centered on the peak response within the response window. The best stimulus was defined as the stimulus evoking the maximum response while marginalizing over other stimulus parameters (for example, the best frequency for pure tones was defined as the frequency evoking the largest intensity-averaged response).

#### Hierarchical clustering

Hierarchical clustering was performed to characterize the diversity in temporal response patterns and response tuning. L5 and L6b ET neurons were pooled together for all clustering analyses, and cluster composition was subsequently compared between subtypes. To characterize temporal response patterns **(Figure 3)**, mean PSTHs averaged across all stimuli and trials were used. Prior to clustering, each neuron’s mean PSTH was rescaled to the range [0, 1]. Pairwise Euclidean distances between rescaled PSTHs were computed across all neurons. The optimal number of clusters was determined by evaluating the Silhouette criterion over a range of 2 to 50 candidate cluster counts (MATLAB function `evalclusters`), with the final count selected at the elbow of the silhouette profile to balance cluster compactness against over-segmentation. Neurons were then assigned to clusters using Ward’s linkage (MATLAB `linkage` and `cluster` functions), sorted by cluster membership, and visualized as ordered PSTH heatmaps. The proportion of L5 and L6b ET neurons in each cluster was quantified and compared.

For pure tone response tuning characterization, hierarchical clustering of frequency-averaged (intensity tuning) and intensity-averaged (frequency tuning) responses was performed as described above **(Figure 4)**. Intensity tuning clusters were reclassified as monotonically increasing, monotonically decreasing, or non-monotonic based on the direction of response change across intensity levels. A monotonic classification required a consistent increase or decrease across at least 4 of 6 intensity levels; remaining clusters were classified as non-monotonic. Frequency tuning clusters were classified as single-peaked (one dominant best frequency region), multi-peaked (two or more discrete response peaks across non-adjacent frequency channels), or complex (no identifiable best frequency region, with broadly distributed responses). Complex profiles were distinguished from multi-peaked profiles by the absence of discrete spectral peaks.

Likewise, sAM noise tuning profiles were characterized by clustering modulation-frequency-averaged responses (depth tuning) and modulation-depth-averaged responses (frequency tuning) independently, as described for pure tones **(Figure 5)**. Depth tuning clusters were classified into four categories: increasing (response grows monotonically with modulation depth), decreasing (response declines monotonically with depth), quadratic (U-shaped profile, with weakest or strongest responses at intermediate depths), or complex (irregular profiles not conforming to any of the above). Modulation frequency tuning clusters were classified as lowpass (strongest responses at low sAM rates), highpass (strongest responses at high sAM rates), bandpass (peak response at intermediate sAM rates), or broadly responsive (no systematic tuning to sAM frequency).

Ripple tuning characterization **(Figure 6)** followed the same clustering approach. Temporal modulation tuning curves were derived by averaging each neuron’s RTF across spectral modulation values, and spectral modulation tuning curves by averaging across temporal modulation values. Both were then clustered as described above. For all stimulus classes, the distribution of tuning categories was compared between L5 and L6b ET neurons.

#### Sparseness and reliability

Sparseness was computed for all pure-tone-responsive neurons. For each responsive neuron (excited and suppressed), the mean response was computed for each stimulus within the response window, resulting in a *N*-element response vector, where *N* is the number of pure tones presented. The resulting response vector was rescaled to the range [0, 1]. The sparseness index (*S_L_*) was computed as [8, 43, 44]:

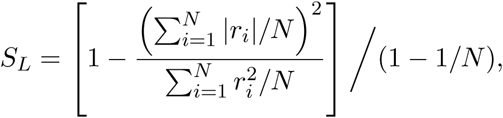

where *r_i_* is the normalized mean response to the *i*-th stimulus, and *N* is the number of presented stimuli.

Single-neuron reliability was computed as the ratio of the variance of the mean response across stimulus conditions (tuning curve variance) to the total variance of single-trial responses pooled across all stimuli (trial variance) [80]:

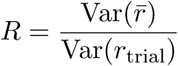

where *r̅* is the mean response across trials for each stimulus condition and *r*_trial_ is the single-trial response. To assess whether reliability differences between L5 and L6b ET neurons were uniform across sound levels or emerged preferentially at specific intensities, we computed reliability indices stratified by sound level. The same reliability metric was applied to sAM noise and spectrotemporal ripple responses.

#### Noise correlations

Noise correlations, also known as spike count correlations, were calculated as the Pearson correlation coefficient of the mean-subtracted trial responses between a pair of simultaneously recorded neurons driven by the same stimulus [51]. For each neuron, the trial response was defined as the mean of the deconvolved spike signal within the response window. The mean response across trials was then computed separately for each stimulus and subtracted from individual trial responses, yielding mean-subtracted residuals that capture stimulus-independent variability. These residuals were concatenated across all stimulus conditions for each neuron in a given field of view, producing a [trials × neurons] matrix from which pairwise Pearson correlation coefficients were computed. To characterize the spatial dependence of noise correlations, intersomatic distances between neuron pairs were calculated as the Euclidean distance between somatic centroid coordinates obtained from Suite2p. For visualization and group comparisons, correlation coefficients were binned by intersomatic distance into 100 *µ*m intervals from 0 to 500 *µ*m, with an additional bin for distances *>* 500 *µ*m. To quantify the rate of spatial decay, an exponential decay function was fitted to the pairwise correlation as a function of intersomatic distance using nonlinear least squares (using MATLAB function `lsqcurvefit`). The decay constant *λ* was extracted per imaging FOV and compared between L5 and L6b ET neurons.

### Statistical Analysis

All statistical analyses were performed in MATLAB (MathWorks) or GraphPad Prism (v10.4.1; GraphPad Software). Data are reported as mean ± s.e.m. unless otherwise stated. Non-parametric statistical tests were used where data samples did not meet the assumptions of parametric statistical tests. All tests were two-tailed, significance was set at *α* = 0.05, and exact *p*-values are reported in the respective Results sections. Significance on figures denoted as follows: * *p <* 0.05, ** *p <* 0.01, *** *p <* 0.0001.

## Author Contributions

MG and RSW conceptualized all experiments. MG conducted all surgical procedures. MG collected and analyzed all data. MG and RSW prepared the figures and wrote the manuscript.

## Acknowledgements

We thank current and former members of the Williamson Lab for helpful feedback and discussions and assistance with animal care. This work was supported by NIH/NIDCD grants R21DC018327 and R01DC020459, and the Klingenstein-Simons Fellowship in Neuroscience to RSW.

